# A Robust, Compact and Diverse Population Code for Competing Sounds in Auditory Cortex

**DOI:** 10.1101/2022.10.24.513560

**Authors:** Jian Carlo Nocon, Jake Witter, Conor Houghton, Howard Gritton, Xue Han, Kamal Sen

**Affiliations:** Neurophotonics Center, Boston University, Boston, Massachusetts, United States of America 02215; Center for Systems Neuroscience, Boston University, Boston, Massachusetts, United States of America, 02215; Hearing Research Center, Boston University, Boston, Massachusetts, United States of America, 02215; Department of Biomedical Engineering, Boston University, Boston, Massachusetts, United States of America, 02215; Department of Comparative Biosciences, University of Illinois, Urbana, Illinois, United States of America, 61820; Department of Bioengineering, University of Illinois, Urbana, Illinois, United States of America, 61820; Department of Computer Science, University of Bristol, England, United Kingdom, BS8 1UB

## Abstract

Cortical circuits encoding sensory information consist of populations of neurons, yet how information aggregates via pooling individual cells remains poorly understood. Such pooling may be particularly important in noisy settings where single neuron encoding is degraded. One example is the cocktail party problem, with competing sounds from multiple spatial locations. How populations of neurons in auditory cortex (ACx) code competing sounds have not been previously investigated. Here, we apply a novel information theoretic approach to estimate information in populations of neurons in ACx about competing sounds from multiple spatial locations, including both summed population (SP) and labeled line (LL) codes. We find that a small subset of neurons is sufficient to nearly maximize mutual information over different spatial configurations, with the LL code outperforming the SP code, and approaching information levels attained without noise. Moreover, with a LL code, units with diverse spatial responses, including both regular and narrow-spiking units, constitute the best pool. Finally, information in the population increases with spatial separation between target and masker, in correspondence with behavioral results on spatial release from masking in human and animals. Taken together, our results reveal that a compact and diverse population of neurons in ACx provide a robust code for competing sounds from different spatial locations.

## Introduction

A central, surprising finding of systems neuroscience is that the discrimination performance of single cortical neurons can match behavior (Britten et al., 1992; Wang et al., 2007). However, an outstanding question is whether single neurons can withstand highly noisy settings, and whether population coding can improve discrimination performance in such settings. An important example of a noisy setting is the cocktail party problem (CPP), where competing sounds originate from different spatial locations (McDermott, 2009). Such settings are highly challenging for a variety of populations with impairments, e.g., ADHD, autism and hearing impairment, assistive devices, and for speech recognition technology, e.g., SIRI and Alexa. Understanding how information is coded in such noisy settings by single neurons and how it aggregates with population coding may illuminate better strategies for treatments in impaired populations, as well as improvements in assistive devices and speech recognition in noise.

Previous studies in songbirds quantified the discrimination performance of single neurons finding degraded discrimination of target sounds in the presence of a competing masker at the same location (Narayan et al., 2007), with high performance levels when the target and masker are spatially separated (Maddox et al., 2012). In this case, the best single neuron code may suffice to support behavior. However, a more recent study in the mouse ACx with a similar experimental design found that discrimination performance of single neurons in the presence of competing sounds is significantly degraded in the presence of competing sounds (Nocon et al., 2022). In this case, population coding may be necessary to support behavior.

Recent studies in ACx have investigated population coding of natural sounds (Ince et al., 2013), and dynamic amplitude modulated sounds (Downer et al., 2021). However, population coding of competing sounds has yet to be investigated, motivating the following questions: Can population coding improve the representation of competing sounds? How do different coding schemes compare? What is the size and nature of the best population under such schemes? Here, we investigate these questions in mouse ACx by applying a novel information theoretic approach. We examine these questions using two population coding schemes: the summed population (SP) code, where response origins are irrelevant to coding, and the labeled line (LL) code, where response origin is maintained (Aronov et al., 2003).

Many information theoretic approaches require binning spike trains at a certain temporal resolution (Ince et al., 2013), whereas the relevant temporal resolution in cortex is unknown. Other approaches have employed spike distance metrics which are free of a choice of temporal resolution (Satuvuori and Kreuz, 2018; Satuvuori et al., 2018). However, spike distance-based approaches have typically been used in conjunction with a specific classifier (Narayan et al., 2007; Wang et al., 2007; Maddox et al., 2012; Nocon et al., 2022). It is unclear how to choose a classifier that best corresponds to cortical computations, or how the choice of classifier influences estimates of discrimination performance. Here we combine the strengths of a time-scale free distance measure (Satuvuori and Kreuz, 2018; Satuvuori et al., 2018), and a recently proposed classifier-free information-theoretic-approach (Houghton, 2015, 2019) to probe population coding of competing sounds in mouse ACx. We find that population coding achieves near maximal information levels with a surprisingly compact population of neurons in both the SP and LL schemes. Furthermore, the LL scheme outperforms the SP scheme, using a population of neurons with diverse spatial responses and cell types, greatly improving information available from single neurons, and approaching information levels of “clean” stimuli without competing noise. Finally, information available in population increases with spatial separation between competing sounds, matching spatial release from masking observed at the behavioral level in animals and humans. Our results reveal a robust, compact, and diverse population code for competing sounds in ACx.

## Materials and Methods

### Dataset

Our dataset consisted of previously collected responses to spatially distributed sound mixtures in right hemisphere of auditory cortex (ACx) from 9 mice (Nocon et al., 2022). All procedures involving animals were approved by the Boston University Institutional Animal Care and Use Committee (IACUC). Subjects consisted of both male and female offspring 8-12 weeks old on the day of recording. During recordings, target and masker stimuli were played from four speakers in space: two contralateral (90º, 45º) to right hemisphere, one at center (0º), and one ipsilateral (∼90º). Target stimuli consisted of white noise modulated in time by human speech-shaped envelopes taken from recordings of a speech corpus (Rothauser, 1969), while maskers consisted of 10 unique tokens of unmodulated white noise. Our past analysis of this dataset was restricted to coding from single units (n = 23). Here, we analyze how coding from populations of neurons facilitate mutual information of stimulus identity during complex scene analysis.

### Population searches using summed population and labeled line hypotheses

Given a full population of N encoding cells X = [x_1_, x_2_, …, x_N_], we wanted to determine the subset within X that best encodes stimulus information, which we define as K_opt_ with size n < N. For each spatial grid configuration, searches for K_opt_ were carried out using either the summed population (SP) or labeled line (LL) approach. The SP code hypothesizes that mutual information is optimized by pooling various neurons to create a single, population-wide response. Specifically, the single response is a union of all spike times from each individual train, and coincident spike times from multiple units are only counted once (Satuvuori and Kreuz, 2018). Meanwhile, the LL code hypothesizes that stimulus features are best decoded on a neuron-by-neuron basis. In this approach, responses from different units were concatenated in time to create a response whose length is the product of the trial length and the number of units.

For all population searches, a bottom-up forward selection algorithm from (Satuvuori et al., 2018) was used. The algorithm first starts with the single neuron that yields the best mutual information and then builds up the population by adding all remaining units at each step, based on the resulting mutual information (Figure 1C). To account for unnecessary additions to K_opt_ due to plateaus in mutual information values, we took the final value of n as the lowest number of neurons that reached 90% of the maximum MI across all subset sizes.

**Figure 1.**
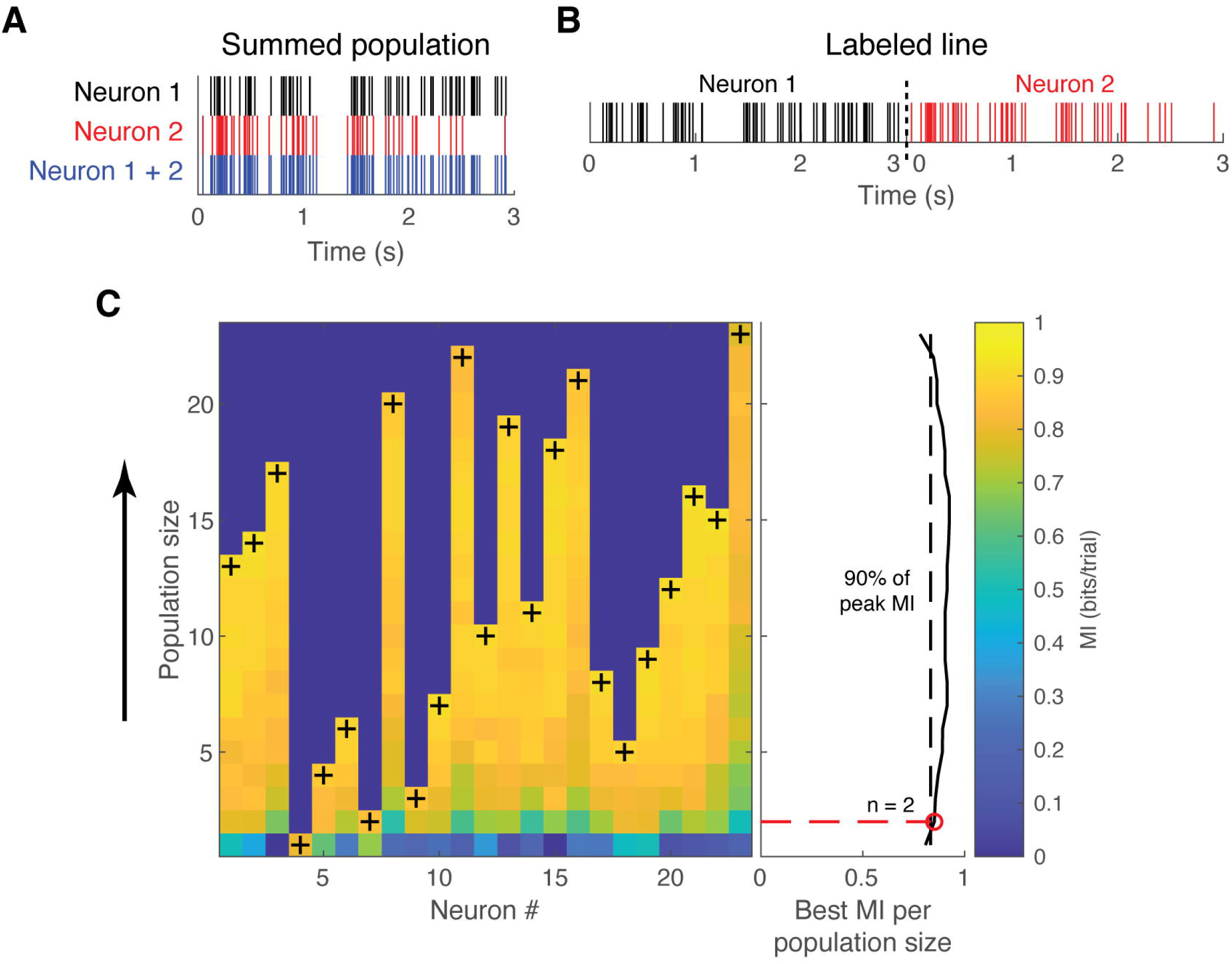
Population approaches and forward selection algorithm. **(A)** Two spike trains in the summed population approach. Spike trains that occur during the same trial are pooled, resulting in a single neural response. **(B)** Example of labeled line approach. The same spike trains in (A) are concatenated in time to create a response whose length is the product of the trial time and the population size. **(C)** Forward selection algorithm. The matrix shows all MI values for each population size, starting with single units at the bottom row. In each row, “+” represents the unit that complements the current population of all prior “+” units. The forward selection search continues by adding each “+” until all neurons have been added. The line plot shows the best MI per population size. To estimate the minimum number of neurons needed to reach optimal MI, we used a threshold at 90% of the maximum MI across all populations (black dashed line). The population that first crosses this threshold was deemed as the optimal neural subpopulation K_opt_ with size n (red open circle and dashed line). Color bar shows the color scale for the mutual information values in the matrix.

### Calculating mutual information using spike train distances

To calculate the mutual information of target identity from spike trains, we utilized an estimator on spike train distances. SPIKE-distance (Kreuz et al., 2013; Satuvuori et al., 2018) calculates the dissimilarity between two spike trains based on differences in spike timing and instantaneous firing rate without additional parameters. For one spike train in a pair, the instantaneous spike timing difference at time t is:

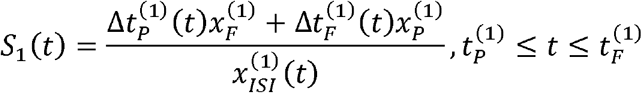

Where Δt_P_ represents the distance between the preceding spike from train 1 (t_p_^(1)^) and the nearest spike from train 2, Δt_F_ represents the distance between the following spike from train 1 (t_F_^(1)^) and the nearest spike from train 2, x_F_ is the absolute difference between t and (t_F_^(1)^), and x_P_ is the absolute difference between t and (t_p_^(1)^). To calculate S_2_ (t), the spike timing difference from the view of the other train, all spike times and ISIs are replaced with the relevant values in train 2. The pairwise instantaneous difference between the two trains is calculated as:

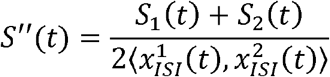

Finally, S_1_(t) and S_2_(t) are locally weighted by their instantaneous spike rates to account for differences in firing rate, and the total distance between the two spike trains is the integral of the profile:

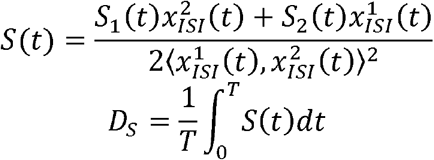

For each single unit, SPIKE-distance matrices were separately calculated for each configuration of target and masker location, with half of the entries representing distances between responses to different target identities and the other half representing distances between responses to the same target.

From these distance matrices, mutual information of stimulus identity was calculated using a Kozachenko-Leonenko estimator (Kozachenko and Leonenko, 1987; Kraskov et al., 2004; Houghton, 2015, 2019), which estimates mutual information on a metric space. It is derived without any reference to a co-ordinate structure, something the space of spike trains lacks. Essentially, it estimates mutual information by approximating probability densities using probabilities

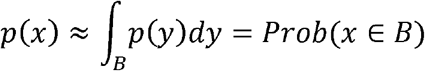

for a region B. The probability Prob(x ∈ B) is then estimated by counting the number of data points in B. In this approach, the distance metric is used to define the small region and the estimator itself is a sort of nearest-neighbor formula, requiring us to look at which data points are near to each other. Calculating mutual information typically requires a lot of data, but this formula appears to work well even when there are not many data points.

## Results

### Description of experimental data and population searches

Using the 23 single units that exhibited at least one “hotspot” of high neural discriminability in response to spatially distributed auditory stimuli from our previously collected dataset (Nocon et al., 2022), we determined the neural subpopulation that gives the highest stimulus identity mutual information (MI) at each spatial grid configuration of target and masker location. To reduce the effect of time resolution on analysis and to avoid the under-sampling problems associated with classical estimators of MI, MI was estimated using spike train distances (Kraskov et al., 2004; Houghton, 2015, 2019). We calculated spike train distances using the SPIKE-distance metric, which measures the difference in local firing rate and spike timing between trains without the need for an additional parameter (Kreuz et al., 2013). We explored two hypotheses of the population code: the summed population (SP, Figure 1A), where spike trains from different neurons are pooled; and the labeled line (LL, Figure 1B), where spike trains are concatenated in time to preserve unit identity. In both approaches, we utilized a forward search algorithm (Satuvuori and Kreuz, 2018), where neurons are added to the population based on how well they complement the mutual information of the current set of neurons (Figure 1C). Because population searches in the labeled line case exhibited plateaus in mutual information as neurons are added to the population, we estimated the minimum number of neurons needed to reach optimal MI. This minimum number was defined as the smallest subpopulation of neurons whose MI was above 90% of the maximum value found across all values of n in the forward search for both population codes.

### Upper envelopes of mutual information

Figure 2Ai shows the upper envelope of optimal MI for the summed population code, while Figure 2Aii shows the upper envelope for the labeled line code. The labeled line search had larger MI values than the summed population approach in almost all configurations, except when the target was at 0º and the masker at 45º. In both cases, a diversity of neuron identities and layer locations was found to contribute to the upper envelope across all configurations, especially in configurations where the target was located at 0º. When comparing how both approaches differed from the best single-unit MI values (bottom-most dashed lines), we found that the labeled line code improved stimulus information coding across all configurations, while the single-unit MI was optimal at some configurations for the summed population code.

**Figure 2.**
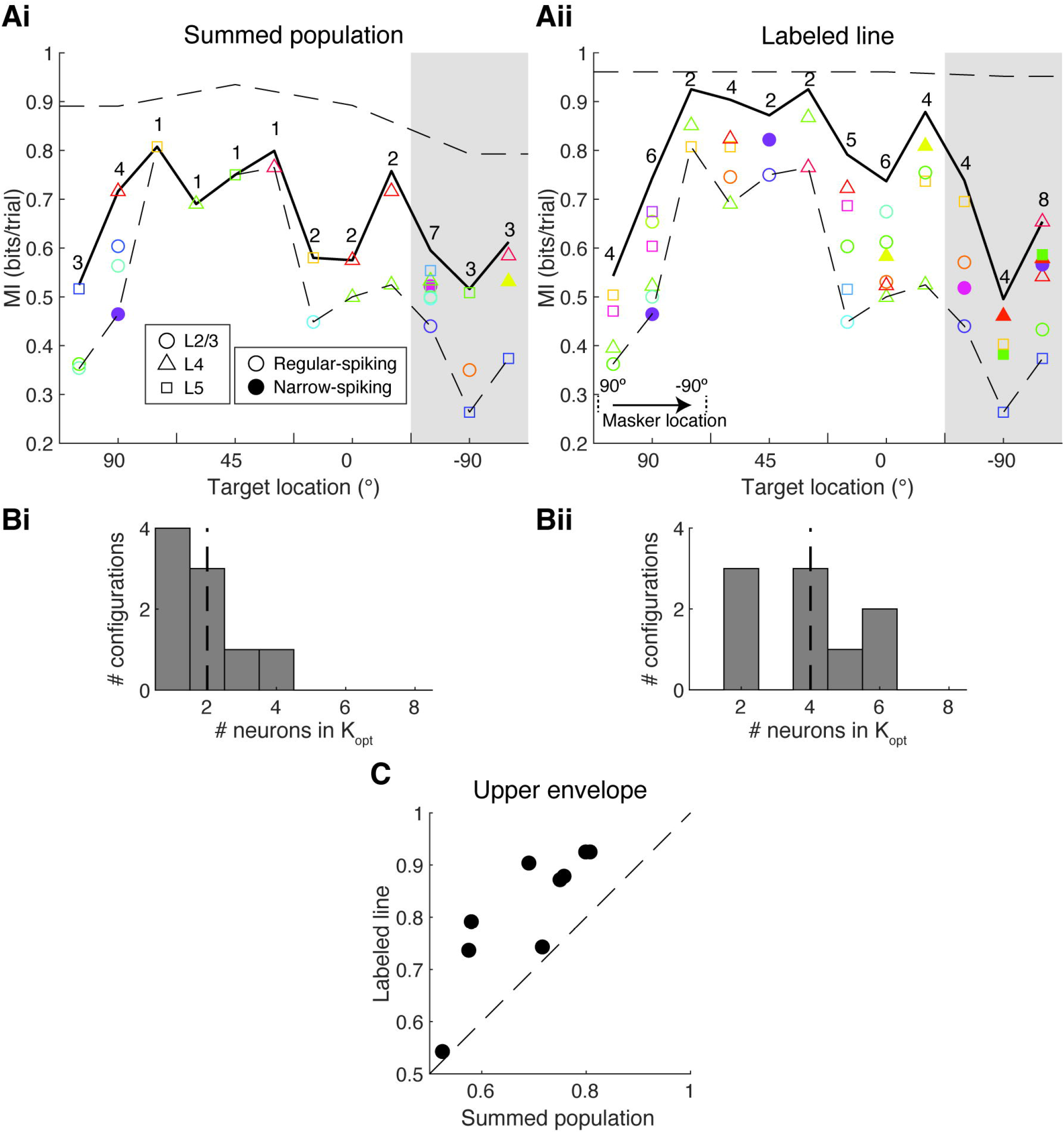
Upper envelopes of mutual information. **(A)** Upper envelope of MI for the summed population **(Ai)** and labeled line **(Aii)** approaches at all spatial grid configurations. Shaded gray region denotes configurations where the target stimulus was ipsilateral (∼90º) to the recorded auditory cortex hemisphere. Bold x-ticks at the bottom separate the plot into sections based on the target location; within each section, the masker location is ordered from contralateral (90º) to ipsilateral (∼90º), as shown by the arrow bounded with dashed lines in bottom-left of Ai. Solid black line represents MI at masked configurations where the target and masker are spatially separated, or non-collocated. Top-most dashed line represents MI at clean configurations. The bottom-most dashed line represents MI from the best single unit (n = 1) at the non-collocated masked configurations. Each separated configuration shows n, the number of neurons in K_opt_. All units included in K_opt_ are shown as markers, with shape representing the cortical layer and color representing the identity of the unit; open markers represent regular-spiking neurons while filled markers represent narrow-spiking units. For some configurations (SP configurations at -90º in SP, all configurations in LL), the highest incremental MI is lower than the upper envelope due to our estimations of K_opt_. **(B)** Histograms of the number of neurons in K_opt_ for all non-collocated, non-ipsilateral target configurations for the summed population **(Bi)** and labeled line **(Bii)** approaches, with the dashed lines representing the median number of neurons per configuration. **(C)** Scatter plot comparing the upper envelope of MI between both approaches at all non-collocated, non-ipsilateral target configurations, with dashed line representing unity.

### Number of neurons in optimal subpopulations and spatial separation between target and masker

For the rest of our analysis, we focused the rest of our analysis on all configurations where 1) the target location is either contralaterally or centrally located relative to the recorded auditory cortex, and 2) the masker stimulus is spatially separated from the target. When we compared the number of neurons that compose the MI upper envelope, we found that the labeled line code had larger subpopulation sizes than the summed population code. Indeed, the median number of neurons in K_opt_ from summed population searches was 2 (Figure 2Bi), while the median number from labeled line searches was 4 (Figure 2Bii). In total, 11 unique units were required to optimally code MI across all optimal subpopulations during SP, while 15 unique units were required during LL. When directly comparing the two codes, we found that the labeled line approach resulted in mutual information values that outperformed those from the summed population approach (Figure 2C). These results, along with the finding that optimal subpopulations in the labeled line codes are larger than those in summed populations, are consistent with (Ince et al., 2013), which found that forward searches for pooled spike trains showed optimal information values at smaller population sizes.

Finally, when we plotted the MIs of separated masked configurations of interest versus the amount of spatial separation between target and masker, we found that MI increased with spatial release for both population approaches (Figure 3). In both cases, there was a significant correlation between the spatial separation of target and masker and stimulus mutual information, which is consistent with findings on the effects of spatial unmasking of target stimuli in humans (Best et al., 2005) and songbirds (Maddox et al., 2012).

**Figure 3.**
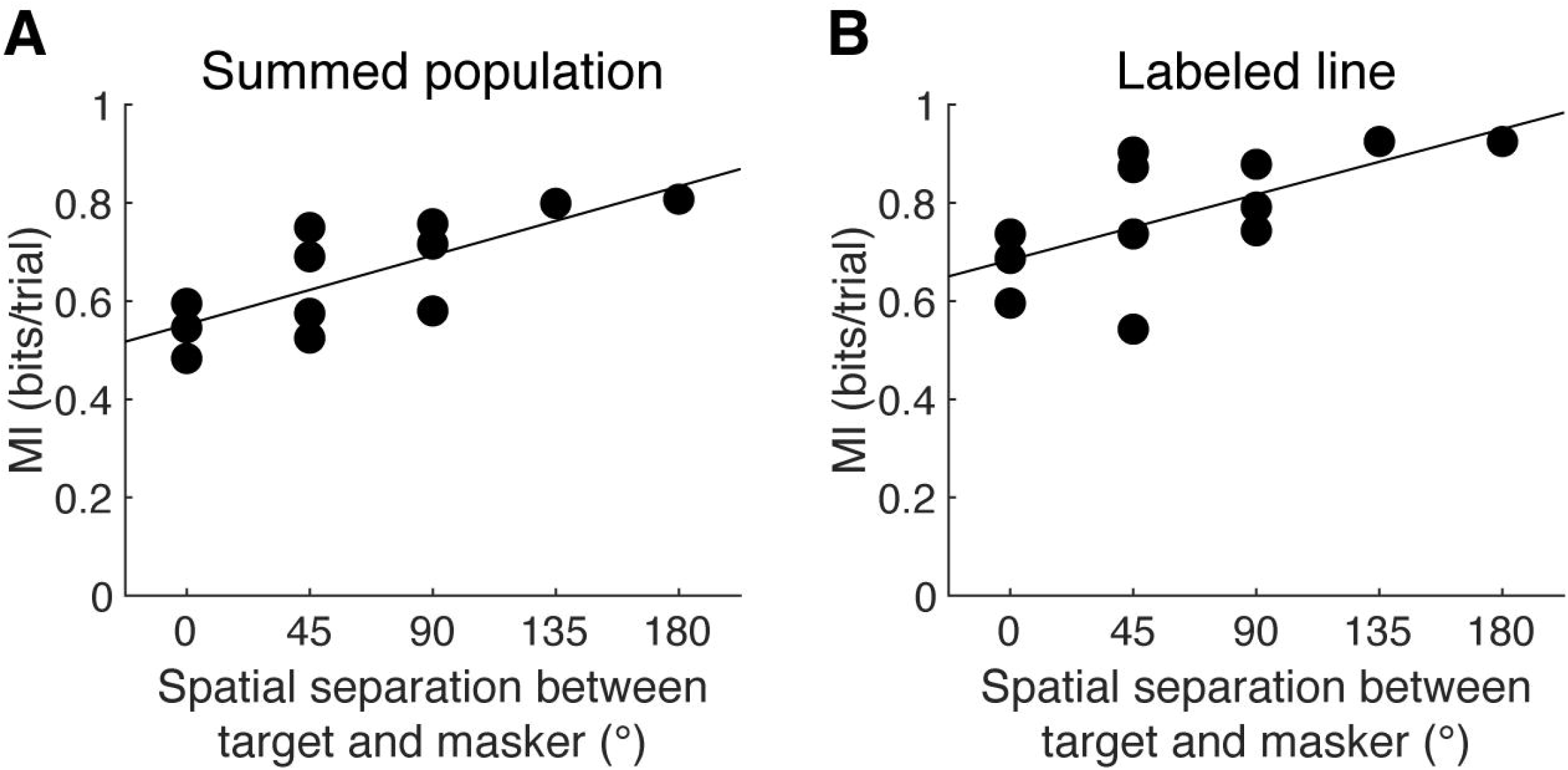
Spatial release from masking versus mutual information. Spatial separation between target and masker versus MI with linear regression lines for both approaches, excluding all ipsilateral target configurations. For both approaches, there was a significant correlation between spatial release and stimulus mutual information (summed population: r = 0.77, p = 0.0036; labeled line: r = 0.65, p = 0.0229).

## Discussion

### Bin-less estimation of mutual information

Previous studies on population coding of auditory stimuli have systematically varied bin sizes on spike times (Downer et al., 2021) or stimulus epochs (Ince et al., 2013), which affect calculations of mutual information provided by spiking events. Here, we present a pipeline for the bin-less estimation of mutual information which involves the use of spike train distance, which are based on differences or similarities between responses without the use of a coordinate system (Houghton, 2015). For this present study, we utilized a time scale-free spike distance metric where the distance between two trains is based on differences in spike timing and local rate (Kreuz et al., 2013; Satuvuori and Kreuz, 2018). The combination of a time scale-free distance metric and estimator decreases effects that firing rate would have on mutual information results and avoids the need to determine time scales that reflect the neural population of interest (Victor, 2002).

### Comparisons between SP and LL codes

In this study, we used two different population coding schemes to determine how stimulus information is optimally coded within mouse ACx. The key difference between these two approaches is whether neuron identity is maintained: in SP, responses are collapsed across time, resulting in a single response. Because of this, SP coding has a destructive effect on the temporal features of individual neural responses, which we previously found to be especially important for high neural discrimination of dynamic stimuli (Nocon et al., 2022). Indeed, for the configurations with contralateral target locations, the median optimal population size was 2 during summed population coding. In this approach, only a few neurons were required to span the entire spatial grid of target and masker location configurations.

In contrast, individual neuron identities and temporal response features were kept intact during the labeled line code. This approach to coding yielded higher optimal MI values at all non-ipsilateral target configurations and improved upon the best MI from single units. Of note is that the optimal subpopulation features both regular and narrow-spiking cells for some configurations (Figure S1), which points to the formation of cortical circuits featuring both types of cells that contribute to high discriminability between target identities

When comparing the upper envelopes from both population codes, we found that the configuration where the target was at 0º and the masker was at 45º not only exhibited similar optimal values, but both approaches required more neurons to approach the upper envelope than other configurations. Interestingly, despite having the same spatial separation between target and masker, the configuration with target at 90º and masker at 45º required less neurons to reach similar optimal MI values. We also found that both configurations share a similar best single unit MI. One possible explanation for this result comes from the contralateral bias in cortex (Woods et al., 2006; Lee and Middlebrooks, 2013): based on the MI upper envelopes in both codes, the population is biased towards target locations at 45º, where the target is not completely contralateral. It is likely that some of the recorded cells within our dataset exhibit a less contralaterally-shifted tuning curve which arises in the population codes.

Our analysis did not include the ipsilateral target configurations due to the contralateral bias present in ACx. Indeed, for most of these configurations, optimal MI was lower than that at other configurations where the target was contralaterally located in both codes. Of interest is the comparison between SP and LL MI at the configuration with target at -90º and masker at 45º, where MI is greater in the summed population code than in the labeled line code. However, the optimal subpopulation size in the SP code includes almost half of the single units used for population coding analysis. It is likely that some cells within that subpopulation encode different temporal features of each target stimulus. We also did not include collocated responses when looking at the number of neurons in the optimal subpopulations, as the degraded response from the competition between target and masker would result in overestimation of the number of neurons needed for optimal MI.

### Spatial release from masking increases stimulus mutual information

Previous studies within cortical responses in songbirds have found that neural discriminability increases with spatial separation between target and masking stimuli. Across both approaches, we found that spatial unmasking of target stimuli increased the optimal stimulus mutual information. These findings are consistent with similar results from behavioral experiments involving both speech and non-speech stimuli (Saberi et al., 1991; Best et al., 2005). It is possible that the labeled line code better reflects the behavioral sensitivity to spatial unmasking, as the cortical processing of cognitive decisions are better explained by circuits containing multiple neuron types.

### Limitations of study

One limitation of our analysis is our sampling of ACx. From our previously collected data, we identified a total of 82 single units across 9 mice, all recorded from right hemisphere. Here, we restricted our population to the 23 single units that showed at least one hotspot of high neural discriminability, and our analysis did not include resampling. In comparison, (Downer et al., 2021) utilized 278 single units across 2 subjects, and their analysis involved repeated sampling of subpopulations of 20 neurons each. In contrast, (Ince et al., 2013) utilized 49 neurons across 3 subjects and did not restrict their population to responsive units only for unbiased analysis. Despite the differences in population sizes and sampling methods, our analysis showed that labeled line code quickly approaches saturation values at around a few (∼5) neurons, while summed population codes are maximized when the population is very small or close to 1, both of which agree with these previous studies.

In this study, neural populations were optimized using a forward search approach. We found that a brute force search consisting of all possible neuron populations did not supplement our analysis, as single unit MIs were already for half of the configurations in the summed population code and labeled line-based MI approached ceiling for some configurations with just one iteration of the forward search. While a brute force analysis would have given us the true subset of neurons that compose the upper envelope, we found that the final step of estimating the minimum population size at each configuration provided the best trade-off between accuracy and computational overhead. Additionally, the population of single-unit neurons used for this analysis was restricted to those that showed at least one performance hotspot during complex scene analysis and thus did not include any non-encoding neurons. We were interested in determining if a population of multiple neurons could improve upon single-unit results, which have already been shown to encode the two target stimuli during clean trials where only the target was present.

Finally, mutual information was based on how spikes in a train contribute to the discriminability between two dynamic auditory stimuli. Because of this, we did not expect the summed population results to greatly improve upon single unit results, which we have already found to encode stimulus identity via template-based classifiers (Nocon et al., 2022). Previous studies on population coding have utilized multiple stimuli (Downer et al., 2021) or binned responses in time (Ince et al., 2013), whereas our calculations of SPIKE-distance were based on differences between trains across the entire period of stimulus playback. Future studies on population coding during complex scene analysis could utilize multiple target stimuli or stimuli with both spectral and temporal differences to better determine how each approach optimizes MI.

## Notes

### Competing Interest Statement

The authors have declared no competing interest.

