## Supplemental Figures for "A Robust, Compact and Diverse Population Code for Competing Sounds in Auditory Cortex"

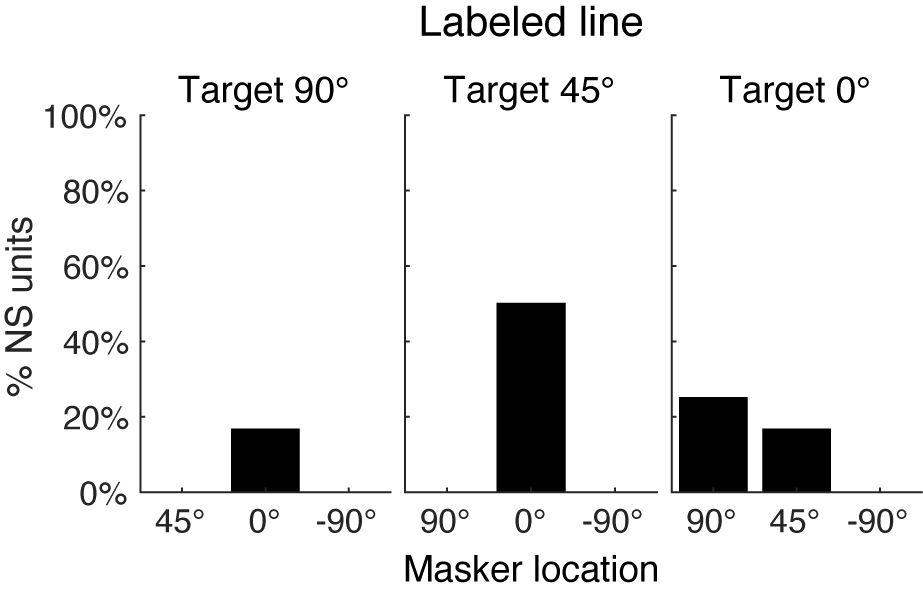

**Figure S1. Narrow-spiking units** **in optimal LL subpopulations.** Proportion of NS units in optimal subpopulation at each non-collocated masked configuration shown in Figure 2Aii.
